# Transcranial or non-transcranial stimulation? Results of a combined TMS-EEG feasibility study

**DOI:** 10.1101/525584

**Authors:** Lutz A. Krawinkel, Julia Forisch, Jan F. Feldheim, Winifried Backhaus, Fanny Quandt, Christian Gerloff

**Affiliations:** Experimental Electrophysiology and Neuroimaging (xENi) Lab, Department of Neurology, University Medical Center Hamburg-Eppendorf, Martinistraße 52, 20246 Hamburg, Germany

## Abstract

**Background:** The combination of Transcranial Magnetic Stimulation (TMS) and Electroencephalography (EEG) provides the possibility to record neurophysiological responses to non-invasive brain stimulation. Previous studies found reproducible time-averaged potentials evoked by TMS (TEPs). Brain electric responses to TMS, including some components of TEPs, might, however, reflect somatosensory and auditory stimulation, rather than direct cortical neuronal responses to the transcranial stimuli. This is of great relevance for interpreting experimental data geared at modulating neuronal networks by TMS. Here we report observations from a feasibility study suggesting that TMS-induced effects on cortical oscillations need to be interpreted with caution.

**Methods:** Subthreshold, monofocal TMS to M1 and PO3 (sham) and bifocal single-pulse TMS was applied to the targets (i) ventral premotor cortex (PMv) and primary motor cortex (M1) and (ii) anterior intraparietal sulcus (aIPS) and M1 using small coils (25mm inner diameter). Besides sham stimulation, we conducted control experiments with a pure auditory and sensory co-stimulation.

**Results:** TMS led to EEG phase synchronization and a pattern of evoked potentials which were comparable with those patterns evoked by sham-stimulation. Moreover, somatosensory stimulation when combined with auditory stimulation mimicked TMS-induced EEG responses. Despite small coils (inner radius 25mm), 3D-neuronavigation analysis showed that aIPS and M1 or PMv and M1 could not precisely be targeted due to their vicinity.

**Conclusions:** TEPs and phase-synchronization of EEG can be affected by TMS-induced non-transcranial somatosensory and auditory components. Simultaneous stimulation of nearby targets like PMv and M1 or M1 and aIPS may be hampered by limited possibilities to position even small TMS coils next to each other.

**Significance Statement:** The present data provide transparency on relevant limitations of a promising neuromodulation approach. While subthreshold TMS can induce resetting and entrainment effects on cortical oscillations, the specificity of these effects needs to be interpreted with caution. Non-transcranial stimulation effects and limitations of coil positioning directly next to each other need to be considered and controlled for.

## 1. Introduction

The combination of Transcranial Magnetic Stimulation (TMS) with Electroencephalography (EEG) offers a broad range of applications: Repetitive TMS (rTMS) modulates ongoing oscillatory brain activity in frequency-specific manner (Plewnia et al., 2008; Thut et al., 2011) and there are long-lasting EEG-aftereffects of theta-burst stimulation (Noh et al., 2012). Moreover, cortical responses to single-pulse and paired-pulse TMS are well characterized (Fecchio et al., 2017; Premoli et al., 2018, 2014; Rogasch et al., 2014; Rosanova et al., 2009) and have been shown to be altered in certain disorders like stroke (Pellicciari et al., 2018), dementia (Ferreri et al., 2016) and especially schizophrenia (Hui et al., 2019; Noda et al., 2018; Rogasch et al., 2015; Tremblay et al., 2019).

However, there is an ongoing debate about the origin of cortical responses to TMS (Belardinelli et al., 2019; Siebner et al., 2019), because both somatosensory and auditory input have been characterized as a source for several components of TMS evoked potentials (TEPs, (Biabani et al., 2019; Conde et al., 2019; Gordon et al., 2018; Herring et al., 2015)). A study by Conde and colleagues systematically evaluated the non-transcranial responses to TMS (Conde et al., 2019) and showed that the characterized potentials (P30, P60, P180, N45 and N100 (Lioumis et al., 2009)) were also provoked by an elaborated sham-stimulation. In this feasibility study, we applied TMS with the initial aim to modulate coupled alpha-band oscillations between aIPS and M1 and between PMv and M1 in a group of healthy, young adults (similar to (Plewnia et al., 2008)).

After the first series of experiments, however, we observed a number of non-specific TMS effects and a limitation due to the close vicinity of the targets in our particular setting. Our findings add to the important ongoing debate regarding TMS-induced EEG responses (Belardinelli et al., 2019; Siebner et al., 2019).

## 2. Methods

### 2.1. Participants

Overall, seven (6 female, 1 male) young, healthy, right-handed participants were included in the study (mean age 26 years, ± 4.1 SD). Generally, single-pulse session was followed by a repetitive TMS session. Resting-state EEG was recorded before and after each TMS session. Four subjects underwent the complete single-pulse session. Out of these four subjects, three subjects completed the repetitive TMS session. One additional subject was included into the control experiment for somatosensory and auditory inputs. Two participants were initially excluded from further testing, because it was not possible to elicit reliable motor evoked potentials (MEPs) through the cap.

Contraindications against TMS were initially checked using a safety questionnaire (Rossi et al., 2011, 2009). All participants gave written informed consent according to the declaration of Helsinki. Study procedures were approved by the local Ethical Committee.

### 2.2. EEG-recordings

EEG was recorded from 60 Ag/AgCl passive electrodes and was referenced to a nose-tip electrode during recordings (BrainAmp MR Plus® amplifier, Brain Products GmbH, Gilching). Positions were arranged in the 10/20 system (easyCAP®, Brain Products GmbH, Gilching, Germany). Recordings were sampled at 5000Hz and amplified in the range of ± 16.384 mV at a resolution of 0.5 µV. To control for MEPs during the experiment, electromyography was measured using a bipolar montage on right first dorsalis interosseous muscle with tendon reference on the index finger (brainAmpExG®, Brain Products GmbH, Gilching, Germany).

### 2.3. TMS-setup

TMS was performed with a Magstim Super Rapid 2® and Magstim Super Rapid 2 Plus® and two small coils with an inner radius of 25mm (P/N 1165-00, connected via an inline inductor, P/N 3467-00, Magstim, Whitland, UK, Fig. 1). TMS was triggered by a Linux-based, custom-made setup, which marked each TMS pulse in the EEG timeline. MEPs for detection of RMT were recorded, amplified and converted using a CED 1902 and a CED micro 1401 (CED, Cambridge, UK). RMT was detected for each stimulator separately. RMT was defined as the lowest TMS intensity which elicited a motor evoked potential (MEP) amplitude >50µV in at least five of ten consecutive trials. Neuronavigation was performed with a Brainsight® system (Rogue Research, Montreal, Canada). Individual high-resolution T1-weighted anatomical images were used to perform 3D-reconstructions (3T Magentom Skyra MRI scanner, Siemens, Erlangen, Germany) in each participant. PMv, aIPS and M1 were marked individually on the basis of critical landmarks (see Fig. 2). Since coils overheated due to high intensities especially during application of rTMS, they were cooled between conditions.

**Figure 1.**
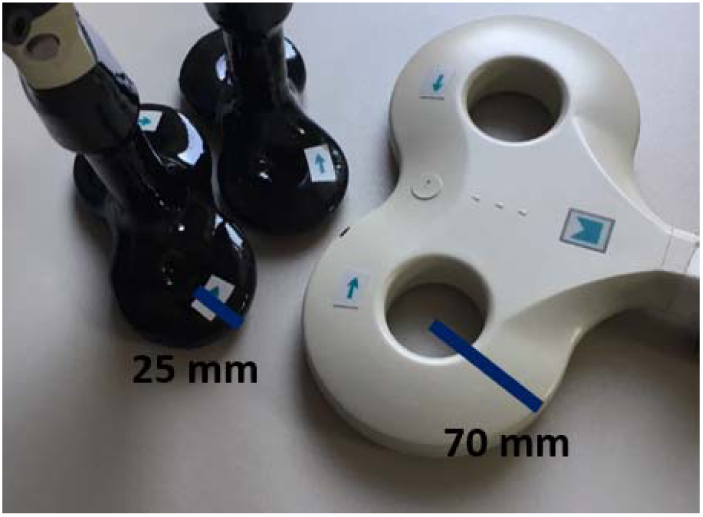
Comparing the Magstim “25mm coils” (left) to a Magstim 70mm coil. TMS with the two “25mm coils” was applied in a shifted fashion to reduce inter-coil-distance.

**Figure 2.**
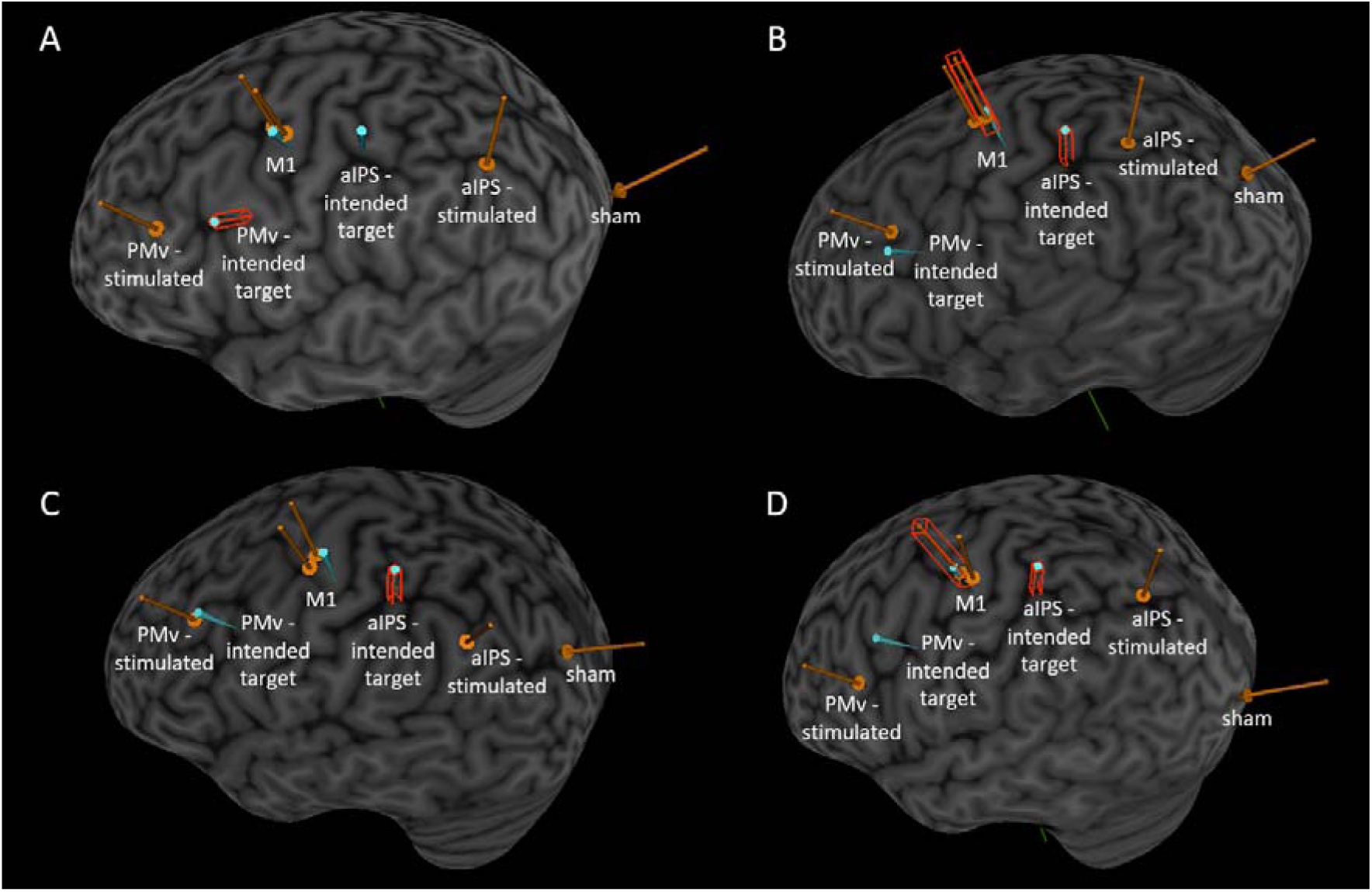
Intended targets (blue) and locations of stimulation (orange) in individual participants (**A-D**). As stimulation was primary adjusted to M1, intended and stimulated locations of the primary motor cortex were closely related. In contrast, intended and stimulated locations of secondary motor areas differed. (Red boxes indicate an active target, topographies were extracted during online navigation-procedure.)

### 2.4. TMS-paradigms and Study Procedure

The experiment took place in an EEG lab with participants seated in an armchair with the right and left arm comfortably in their lap. Participants were instructed to avoid eye blinks and movements. A full-HD monitor (viewangle ±5°, Monitor 24”, Dell, Texas, USA) presented a fixation cross during all trials.

In an initial step, we tested whether it was possible to elicit MEPs in our participants wearing an EEG-cap (easyCAP®, Brain Products GmbH, Gilching, Germany). Subsequently, EEG electrode-positions were registered using an ultrasound localization system (CMS20, Zebris, Isny, Germany) and the individual resting-motor-threshold (RMT) wearing the EEG-cap was measured. Afterwards, the EEG-cap was prepared using abrasive gel in order to keep impedances below 10kOhm. In a fourth step, neuronavigation of M1, PMv and aIPS was performed ensuring correct target locations of the TMS coils relative to individual brain structure. In general, we stimulated the left, dominant hemisphere. Target positions for PMv, M1 and aIPS were marked on the EEG cap. Then the recording session started.

The first experiment comprised four conditions with 90% of RMT both for the singe-pulse (4 participants) and the rTMS session (3 participants): (i) bifocal aIPS-M1 stimulation, (ii) bifocal PMv-M1 stimulation and (iii) monofocal M1 stimulation as a control for M1 effects of stimulation. Application of TMS over electrode position PO3 (10-20 system) served as a (iv) sham condition because we did not expect any responses in the motor system to parieto-occipital stimulation. Stimulation entities (verum/sham) were randomized across participants. Since major findings and limitations of this study arouse after analyzing the single-pulse control experiments, we restrict our analysis to this part of the experiment.

A total of 32 trials was recorded for each of the 4 conditions during the single-pulse session. Trials had an overall length of 8s, with at least 3s pre-stimulus and 3s post-stimulus interval. A jittered interval (0-2s) was added to the pre-stimulus interval, and the remaining time to 2s was added to the post interval. Thus, in total an interval of 2s was added to each trial. The inter-trial interval was 6s.

One participant participated in an extended experimental session, in which several additional control conditions were recorded. Importantly, an auditory stimulation was performed by holding one TMS coil behind the participant’s head (no contact with the skin). The stimulation was applied using the distance from left ear to M1, holding the coil planar and pointing to the participant’s left ear with an angle of 45°. Another condition was a pure sensory, trigeminal stimulation of the left temporal face area being not covered by the EEG cap. For this condition a constant current stimulator (Digitimer, DS7A, Welwyn Garden City, UK, also triggered by the Linux-based-setup) was used, applying the intensity at sensory threshold (3.7mA, 400V max, pulse width of 50µs). A final control condition combined somatosensory and auditory stimulation. The trial length of the adjusted setup was analogue to the original study setup.

### 2.5. Data-Analysis

For EEG phase analysis, raw data were processed into trials from −2 to 2s with respect to the TMS pulse. Second, the TMS artifacts were removed by interpolating an interval of 15ms following each TMS pulse and a 5ms interval around the TMS-recharge artefact (Thut et al., 2011). After trial rejection (for the verum condition 2.5 trials (± 1.3) were rejected, for the sham condition 3.75 trials (± 2.2) were rejected), data were down-sampled to 1000Hz and reverse filtered (8-12Hz and 18-22Hz). One pass reverse filtering was used to avoid a contamination by filter-ringing in the signal following TMS pulse. After filtering, data was shifted by half the filter order to correct for filter-induced offset. Then, data was Hilbert-transformed using filtered data (alpha-band and beta-band) and converted to data containing phase-angles. After that, circular statistics were performed using the function “circ_r” (MATLAB toolbox for circular statistics (Berens, 2009)) resulting in a mean vector, reflecting “directedness” of phase distribution: The closer it is to one, the more concentrated the data sample is around the mean. For statistical analysis “directedness” was averaged over bins of 50ms, except for the period of the TMS-pulse (see below). We computed and plotted “directedness” from −100ms to 1000ms, except for the period of the TMS-pulse: We did not include the period from −20ms to 20ms into our analysis: Due to potential residual artefacts and the reverse filtering, potential disturbances of the filter would smear both into the signal after and especially into the signal before the interpolated TMS-artefact. Statistics were then performed using the Students t-test. Data was Bonferroni-corrected for 22 time-points (bins). For a better visualization data was smoothed using a Savitzky-Golay FIR smoothing filter.

For analysis of TMS-evoked (EEG) potentials (TEPs), the data from 20ms to 2000ms after the TMS-pulse and from 2000ms to 0ms before the TMS pulse was first filtered with a highpass filter (4Hz). After trials with excessive muscle or eye-blink artefacts were rejected (for the verum condition 2.5 trials (± 1.3) were rejected, for the sham condition 3.75 trials (± 2.2) were rejected), data were time-averaged for the period from −40ms to 360ms. Averaged Data is presented as a mean with standard deviations. For TEP-analyze we did not perform statistical analyses due to potentially remaining decay-artefacts and related potential false-positive findings. For a better visualization data was smoothed using a Savitzky-Golay FIR smoothing filter. Single-subject data of the added control experiment were analyzed identically.

In general, the EEG data were analyzed with the FieldTrip package (Oostenveld et al., 2011) on the basis of MATLAB, version 9.2.0.538062 (Mathworks Inc., Massachusetts, USA).

## 3. Results

All participants tolerated the various forms of single-pulse TMS and rTMS well. No severe adverse effects occurred. However, in one subject of this study and in one subject of the pilot-trials to this study, subthreshold TMS at 90% RMT became suprathreshold and elicited MEPs when applied as simultaneous bifocal TMS with two coils positioned directly next to each other. Since subthreshold bifocal stimulation was attempted, these two subjects were excluded from further experiments.

### 3.1. TMS intensities

As described in section 2.1, initially, RMT was detected in each participant. In one participant, it was not possible to elicit stable MEPs (with the coil placed on top of the cap-mounted electrodes). In another participant, it was not possible to elicit any EMG response at all. For any other data shown in this paper, TMS was applied with 90% RMT which corresponded to 81% stimulator output (±7%) on the Super Rapid 2 and 80% stimulator output (±7%) on the Super Rapid 2 plus, respectively.

### 3.2. Quality of neuronavigation

Irrespective of neuronavigation and the already very small coil size (Fig. 1), aIPS and M1 or PMv and M1 could be roughly targeted but the deviation from anatomical target was only acceptable in participants with large head circumference. Figure 2 shows individualized target areas of the four participants (A-D) who completed the single-pulse TMS session. As stimulation was primarily adjusted to M1, intended and stimulated locations of the primary motor cortex were closely related. In contrast, intended and stimulated locations of secondary motor areas (aIPS and PMv) differed.

### 3.3. EEG responses to TMS

Since this study focuses on the modulation of ongoing oscillatory activity, TMS-related local phase locking of 10Hz alpha-band and 20Hz beta-band oscillations, relevant frequency bands of the human motor system (Bönstrup et al., 2018, 2015; Dubovik et al., 2013), was computed as a first step. To comprehend EEG responses to a TMS application with the 25mm coils, EEG during monofocal, single-pulse stimulation of M1 compared to sham stimulation (PO3) was examined in a first step.

Computing the length of the mean vector of phase distributions (here called “directedness”) revealed a clear phase-locking of alpha-band oscillations to the TMS-stimulus at electrode C3, covering the left sensorimotor cortex and peaking in a period from TMS pulse to about 200ms and at about 300ms (Fig. 3A, main panel). Rather unexpectedly, a similar phase-locking pattern at electrode C3 was observed after sham stimulation (Fig. 3A, main panel). Visually, local alpha-band directedness at C3 seemed to be increased after verum stimulation compared to sham stimulation. However, statistical analysis did not reveal significant differences between conditions after correction.

**Figure 3.**
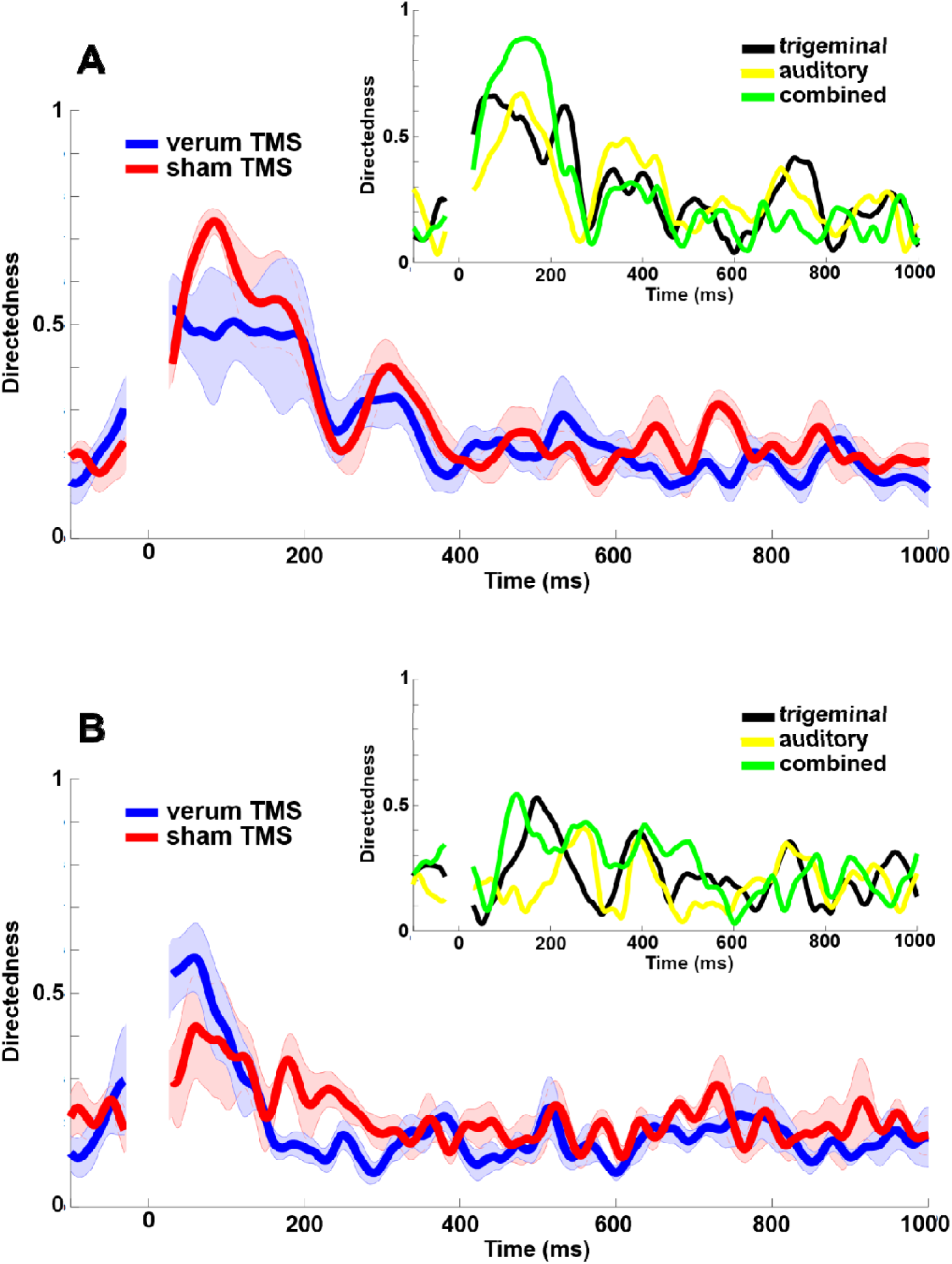
**(A)** Alpha-band and **(B)** beta-band phase properties at electrode level C3. Main panel shows directedness from −100ms to 1000ms with a peak after both verum and sham application of TMS at 0ms. Inlay panel shows directedness after trigeminal, auditory and combined stimulation, indicating a comparable pattern of phase synchronization..

Additionally, we computed directedness at electrode level C3 for beta-band oscillations (Fig 3B, main panel). Generally, phase locking after TMS was less pronounced compared to the alpha-band. However, there was a clear peak after the stimulation. On visual inspection, local beta-band directedness at C3 seemed to be increased after verum stimulation compared to sham stimulation. However, exploratory statistical analysis did not reveal significant differences between conditions for the beta-band.

To further clarify the influence of somatosensory and auditory input induced by subthreshold TMS in our setup (Conde et al., 2019; Herring et al., 2015), we analyzed data of an additional session including somatosensory (trigeminal) and auditory stimulation in a single subject. Both for alpha-band and beta-band oscillations (Fig 3A and 3B, inlay panels) analyses revealed a clear phase-locking to the stimulus. Like after TMS, phase locking was pronounced in the alpha-band compared to the beta band. Generally, the combined auditory and somatosensory stimulation led to the highest increase of directedness, however, the peak at around 300ms to 350 ms was most pronounced after pure auditory stimulation (Fig 3A and 3B, inlay panels).

Apart from phase analyses, we performed the established TEP analysis (Biabani et al., 2019; Conde et al., 2019; Fecchio et al., 2017; Gordon et al., 2018; Lioumis et al., 2009; Rogasch et al., 2015, 2014) to further evaluate effects of our subthreshold TMS-setup. In line with the results of directedness, TEP analysis revealed that both verum-M1-stimulation and sham stimulation elicited the well characterized TEP patterns with positive peaks at around 30ms, 60ms and 180ms (P30, P60, P180) and negative peaks at around 45ms and 100ms (N45, N100 (Lioumis et al., 2009)) at electrode level C3 (Fig 4, main panel). Additionally both stimulation entities revealed a positive peak at around 80ms (Fig 4, main panel).

**Figure 4.**
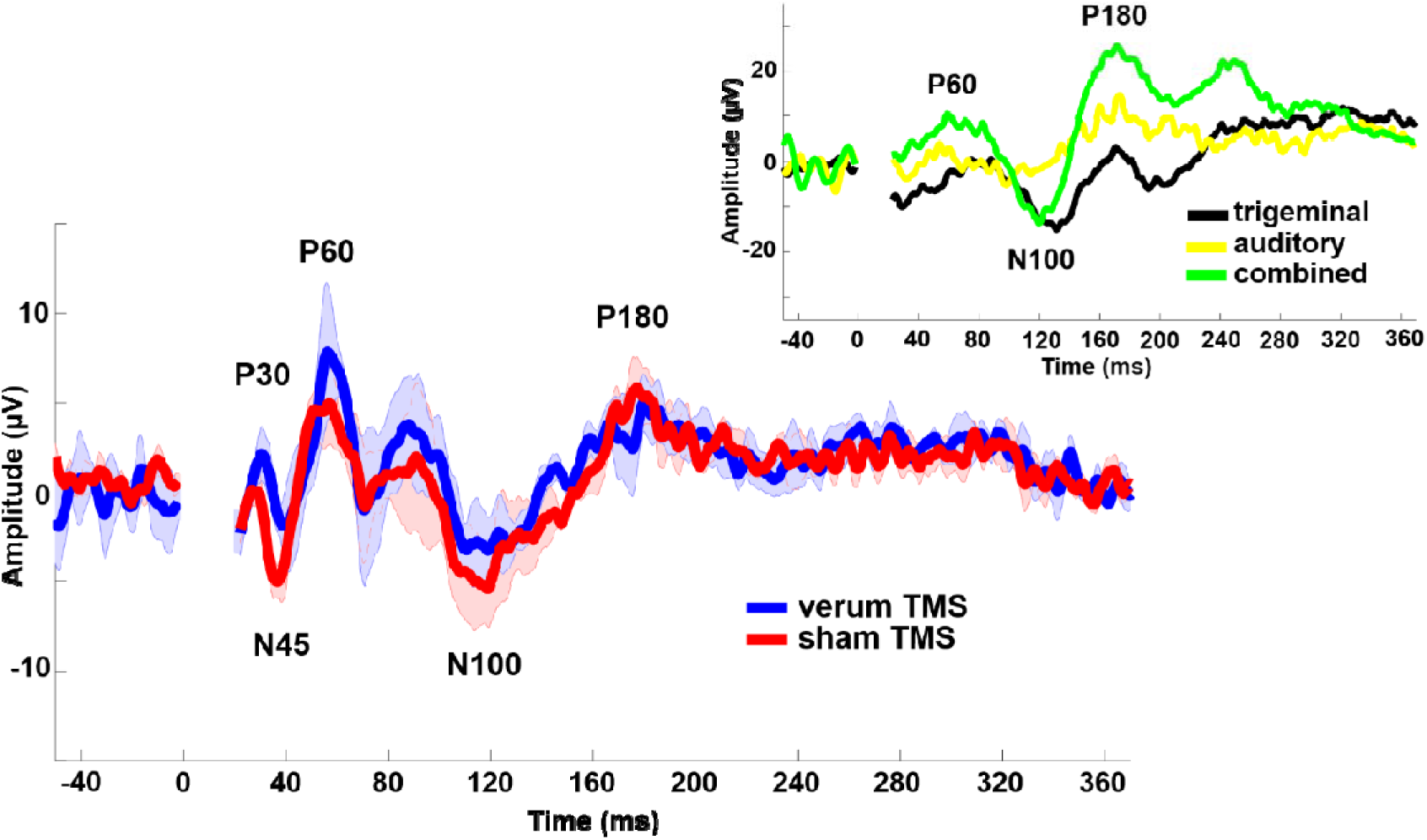
TMS evoked potentials at electrode level C3. TEPs are plotted from −40ms to 360 ms regarding the TMS application at “0ms”. Both verum and stimulation elicited positive peaks at around 30ms, 60ms and 180ms (P30, P60, P180) and negative peaks at around 45ms and 100ms (N45, N100, main panel). Inlay panel shows evoked potentials after trigeminal, auditory and combined stimulation. Especially combined trigeminal and auditory stimulation elicited positive peaks at around 60ms and 180ms (P60, P180) and a negative peak at around 100ms (N100).

Moreover, additional TEP analyses of single-subject data using pure trigeminal and auditory stimulation revealed that especially combined trigeminal and auditory stimulation elicited positive peaks at around 60ms and 180ms (P60, P180) and a negative peak at around 100ms (N100, Fig. 4, inlay panel).

## 4. Discussion

Inducing phase synchrony by non-invasive brain stimulation is a potentially interesting and powerful technique to study and, ideally, enhance neuronal function (Ahn et al., 2019; Romei et al., 2011; Sauseng et al., 2009; Thut et al., 2011; Zaehle et al., 2010), especially because it is conceptually suited to enhance interregional coupling of oscillatory brain activity (Helfrich et al., 2014; Plewnia et al., 2008; Schwab et al., 2019). Based on these considerations, we aimed to establish a feasible setting for entraining key cortical areas known to be engaged in the motor recovery process after stroke (Bönstrup et al., 2018, 2016; Rehme et al., 2011; Schulz et al., 2015). A crucial element of our approach was to detect robust changes of inter-areal coupling metrics (e.g., phase-angles) in EEG (Thut et al., 2011), adjust the stimulation paradigm to optimize these neurophysiological effects, and then enhance the sample size and measure potential effects on motor behavior and recovery after stroke. However, when performing the first series of experiments with subthreshold single-pulse TMS in healthy control participants, we observed technical and physiological effects that finally questioned the application of (r)TMS in this setting.

There turned out to be two key observations that need to be discussed. First, major components of EEG responses to subthreshold TMS were also found after sham stimulation, and could be attributed to the auditory and somatosensory stimulation related to coil discharge, which is in line with some previous findings (Biabani et al., 2019; Conde et al., 2019; Herring et al., 2015). Second, bifocal TMS, even with small 25mm coils, has substantial limitations as to how closely neighboring target areas can be located for precise bifocal stimulation, i.e., aIPS-M1 as used here, or PMv-M1 as used here and previously (Koch et al., 2010).

### (1) Restrictions by closely adjacent target areas and small TMS coils

An important aspect of this observational study was the finding that despite having used very small coils, it was not possible to target M1-PMv or M1-aIPS bifocally as planned in most of the participants (Fig. 2). Only with a very large head circumference these sites can be targeted with low positional error. As a rule, it was not possible to stimulate anterior parts of the IPS together with M1. Only in one subject with a large head, PMv stimulation came close to the intended target if the coil over M1 was positioned first. Our coils had an inner diameter of 25mm. Already with these coils, finding the hotspot and determining the RMT can be challenging. Out of seven young and healthy participants, MEPs, required for measurements of RMT, could only be elicited in five. In elderly participants and stroke patients, the RMT is expected to be higher (Cakar et al., 2016). Coils smaller than 25 mm might theoretically solve the problem of neuronavigation but would probably lack the field strength necessary (and heat up on repetitive stimulation). To our knowledge there are, anyway, currently no CE certified coils commercially available with a smaller diameter than the Magstim® 25mm coils. The problem of low field strength is naturally even greater when TMS is combined with EEG because the electrodes mounted in the cap cause a distance of the coil from the scalp of at least approximately 2mm, rather more. The strength of the magnetic field is maximal close to the coil surface and decreases exponentially with the distance from there (Cohen et al., 1990; de Goede et al., 2018). An interesting way to attenuate this problem might be closed-loop stimulation, through which TMS efficacy might be enhanced by better timing relative to the ongoing oscillatory cortical activity (Bergmann et al., 2012; Kraus et al., 2016; Raco et al., 2016).

Furthermore, with respect to technical innovation, there might be future devices that can offer smaller, cooled coils, e.g., in a cap or helmet setup, which could overcome the spatial restrictions demonstrated in this paper but to our knowledge no such device is CE certified and commercially available as of now.

### (2) Specificity of EEG responses

Both analysis of phase properties and TEP analysis revealed that responses to verum and sham-stimulation were highly similar.

Phase analysis by means of directedness showed that both after verum and after sham application of TMS, phase synchronization increased and patterns where highly comparable (Fig 3A and 3B, main panel). Moreover, the patterns of increased directedness induced by TMS could also be elicited by pure trigeminal and auditory input, and even more so by combined trigeminal and auditory stimulation (Fig 3A and 3B, inlay panels), highly supporting a major influence of non-transcranial mechanisms. The fact, that directedness at around 300-350ms at C3 for the alpha-band (Fig 3A, right box) had the highest level after pure auditory stimulation might underline the role auf auditory stimulation for this peak. However, due to the single subject data, this interpretation remains speculative.

The concept that somatosensory and auditory inputs reflect a major component of EEG responses to TMS (Belardinelli et al., 2019; Biabani et al., 2019; Conde et al., 2019; Gordon et al., 2018; Herring et al., 2015; Siebner et al., 2019) is supported by our TEP analysis. In line with the results of directedness, analysis revealed, that both verum-M1-stimulation and sham stimulation clearly elicited known TEPs with positive peaks at around 30ms, 60ms and 180ms (P30, P60, P180) and negative peaks at around 45ms and 100ms (N45, N100 (Lioumis et al., 2009), Fig. 4, main panel). Additionally, we observed a consistent positive component at around 80ms, which can be also seen in earlier studies (Lioumis et al., 2009). Generally, from an electrophysiological point of view, analogous results of TEP and phase analysis are possibly owing to the fact that phase synchronization is a causal factor for the presence of time-averaged evoked potentials (Makeig et al., 2002).

Our control experiment with pure trigeminal and auditory stimulation supports a major influence of peripheral inputs on TMS-induced responses (Nikouline et al., 1999; Tiitinen et al., 1999). Especially combined trigeminal and auditory stimulation elicited TEPs with a P60, N100 and P180 component (Fig 4, inlay panel). Moreover, figure 3A (inlay panel) clearly shows alpha-band phase synchronization at electrode-level C3 after pure auditory, pure trigeminal and combined stimulation, comparable to the effects after verum stimulation and sham stimulation (figure 3A, main panel). The effects of pure auditory stimulation (no contact to the subject with the coil) underline our interpretation, that volume conduction cannot be the reason for the similarities between the effects of real and sham stimulation. However, after peripheral stimulations, components with a smaller amplitude like P30 or N45 could not be identified (Fig 4, inlay panel). Reasons might be the following: Firstly, small amplitude components might need averaging over multiple subjects. Secondly, a failure to detect these components might be due to the positioning of the peripheral sham-stimulation consisting of a stimulation of the ipsilateral temporal face and an auditory click while the coil was positioned without contact to the bone. In contrast, Conde and colleagues, who could elicit a P30 and N45 with a sham-stimulation, used an elaborated setup with a topographically identical positioning of verum and sham-stimulation. The authors conclude, that early potentials might be, next to early trigeminal input, caused by stimulation parasagittal fibers of the dura mater (Conde et al., 2019).

Though it seems, that non-transcranial trigeminal and auditory inputs have a huge impact on EEG responses to TMS, this study cannot exclude, that there are added or hidden transcranial responses respectively. Supporting the notion, that TEPs reflect transcranial cortical stimulation, previous studies did find differences in TEPs in diseases like schizophrenia being correlated with behavior and being specific for certain brain regions like dorsolateral prefrontal cortex (Hui et al., 2019; Rogasch et al., 2015; Tremblay et al., 2019).

Nevertheless, our analysis clearly pointed out a severe contamination of the EEG with non-transcranial effects. Since this observation is found already in a small sample size, the risk of a contamination in any TMS-EEG has to be regarded as high.

To our understanding, a detailed characterization of very early potentials would be crucial and might bring a benefit to the discussion about the meaning of TEPs, since early potentials cannot be elicited by afferent sensory activity. Since classical double-pulse TMS-paradigms like short intracortical inhibition (SICI, interstimulus interval 1-6ms) or short intracortical facilitation (ICF, interstimulus interval 6-30ms) clearly pointed towards a direct transcranial and transsynaptical stimulation by subthreshold TMS (Rossini et al., 2015), early TEPs might reflect transcranial stimulation.

Generally, according to the data presented here and to recent publications (Conde et al. 2019, Gordon et al. 2018), the following cautions and controls might be considered: (i) an elaborated sham condition including somatosensory and auditory components should be performed at the target of interest. Subjects should be interviewed afterwards to control for a sufficient blinding (Conde et al. 2019, Gordon et al. 2018); (ii) a second control site should be stimulated to prove local-specific transcranial effects (Rosanova et al. 2009); (iii) since early TEPs - before a potential somatosensory or visual re-afference – might reflect transcranial stimulation and are potentially contaminated by TMS-induced electrical and tissue artefacts (Conde et al. 2019, Gordon et al. 2018, Thut et al. 2011), a consensus regarding cleaning methods should be reached.

### Conclusion

Though the impact of our feasibility study is limited due to a small sample size, it underlines the importance of reliable control and sham conditions in TMS studies respectively (Belardinelli et al., 2019; Siebner et al., 2019): In line with previous findings, major components of evoked potentials after TMS might be due to somatosensory and auditory input (Biabani et al., 2019; Conde et al., 2019; Herring et al., 2015). Moreover, our analyses suggest, that oscillatory properties by means of phase synchronization might also be triggered by peripheral inputs and not by a transcranial stimulation.

Furthermore, bifocal TMS, even with small 25mm coils, has substantial refinements as to how closely neighboring target areas can be.

## 5. Acknowledgements

This research was supported by the German Research Foundation: SFB936 “Multi-site communication in the brain”, project C1.

